# *In situ* structure determination of virus capsids imaged within cell nuclei by correlative light and cryo-electron tomography

**DOI:** 10.1101/2020.02.06.936948

**Authors:** Swetha Vijayakrishnan, Marion McElwee, Colin Loney, Frazer Rixon, David Bhella

**Affiliations:** MRC-University of Glasgow Centre for Virus Research, Sir Michael Stoker Building, Garscube Campus, 464 Bearsden Road, Glasgow G61 1QH, Scotland, UK

**Keywords:** Virus, Capsid, Herpesvirus, Cellular imaging, Cryo Electron Microscopy

## Abstract

Cryo electron microscopy (cryo-EM), a key method for structure determination involves imaging purified material embedded in vitreous ice. Images are then computationally processed to obtain three-dimensional structures at atomic resolution. There is increasing interest in extending structural studies by cryo-EM into the cell, where biological structures and processes may be imaged in context. The limited penetrating power of electrons prevents imaging of thick specimens (>500 nm) however. Cryo-sectioning methods employed to overcome this are technically challenging, subject to artefacts or involve specialised equipment of limited availability. Here we describe the first structure of herpesvirus capsids determined by sub-tomogram averaging from nuclei of eukaryotic cells, achieved by cryo-electron tomography (cryo-ET) of re-vitrified cell sections prepared using the Tokuyasu method. Our reconstructions reveal that the capsid associated tegument complex is present on capsids prior to nuclear egress. We show that this approach to cryogenic imaging of cells is suited to both correlative light/electron microscopy and 3D structure determination.

## Introduction

Three-dimensional (3D) information on protein structure is key to our understanding of dynamic processes that occur within a cell or tissue at the molecular level. Electron microscopy (EM) has been a critical tool in the study of cellular ultrastructure. Classical methods for imaging of biological material in the electron microscope involve the use of metal contrasting agents and dehydration of the sample^1^. Such methods limit the preservation and visualisation of fine features. Cryo-EM has recently emerged as a powerful tool in the study of macromolecular structure. Purified protein or nucleoprotein assemblies are embedded in vitreous ice yielding frozen-hydrated specimens that are suited to imaging in the transmission electron microscope. This method produces artefact-free, high-resolution images of biological assemblies in near-native conditions that may be processed to determine high-resolution structures. Cryo-EM has transformed the field of structural biology providing rapid access to high-resolution macromolecular structure without the need to prepare crystals. These studies have however largely been limited to single-particle analysis - the study of purified homogeneous specimens, which may be computationally averaged to yield high-resolution structure data^2^. Many of the most interesting biological questions involve unique and heterogeneous structures that are not amenable to structural averaging methods employed in cryo-EM single-particle techniques. To address this challenge, considerable efforts have been made in the development of cryo electron tomography (Cryo-ET) as a tool to image preparations of heterogeneous assemblies such as enveloped viruses as well as vitrified cells^3-5^. Electron tomography is a means of determining three-dimensional structures of unique entities by recording multiple images of an object in the transmission electron microscope (TEM), as it is rotated, typically through ∼120°. Owing to the limited penetrating power of electrons, cryo-ET is restricted to samples that are less than 500 nm thick, thus for cellular imaging, only the thin edges of cells are accessible. Features in the interior of eukaryotic cells, such as the nucleus, are not readily viewed.

To image the cell interior requires thinning of the area of interest to less than 500 nm. Early attempts to address this problem employed cryogenic microtomy or ‘cryo-sectioning’ of vitreous frozen-hydrated cells. While cryo electron microscopy of vitrified sections (CEMOVIS) avoids chemical fixation and staining, it is technically challenging, limited in sample thickness and subject to several artefacts including compression and crevasses. These restrict the potential of CEMOVIS for high-resolution structural imaging ^6, 7^. More recently progress has been made towards artefact-free cell sectioning through milling of cellular specimens using cryo focused ion beam scanning electron microscopy (cryoFIBSEM)^8^. Frozen-hydrated cells, grown on an EM grid are etched into electron-transparent lamellae, using a gallium ion-beam. Lamellae are then imaged by cryo-ET. When successfully applied cryoFIBSEM provides well preserved views of the cell interior. Focussed ion-beam milling of vitreous specimens is, however, a low throughput method that is prone to failure – lamellae are easily broken or contaminated by frost during transfer from the cryoFIBSEM to the cryo-TEM. Moreover, the method involves specialised and expensive equipment.

Artefact-free or low-artefact cryo-microtomy approaches to preparing cellular material for cryo-ET are therefore desirable. Such approaches have the benefit of using less expensive and more widely available equipment for high-resolution 3D structural imaging of cells and tissues, potentially leading to *in situ* structure determination. Vitrified frozen cell sections (VFS) provide an ideal compromise wherein chemically fixed samples are cut under cryogenic conditions, thawed and then re-vitrified by plunge-freezing for imaging by cryo-EM or cryo-ET ^9, 10^. This technique has several advantages for investigators interested in determining macromolecular structures *in situ*. Many laboratories will have ready access to the necessary equipment; cryo-microtomes are already available in many EM laboratories, or existing microtomes may be modified for low-temperature microtomy. The method is relatively high-throughput; many sections may be prepared at once, with a greater likelihood of successful imaging. To our knowledge however, use of this method has thus far been confined to 3D imaging of tissue specimens ^10-13^. Sub-tomogram averaging has not been hitherto attempted.

While cryo-ET potentially allows imaging of macromolecular structures in 3D within intact cells, it remains challenging to identify features of interest within the crowded and complex environment of the cell. Such studies are therefore liable to be both challenging and time-consuming especially when structures under investigation are smaller, involved in dynamic or rare events or are not well characterized. To overcome these difficulties methods have been developed that allow investigators to precisely locate features of interest prior to recording tomograms. This is achieved through fluorescence microscopy of cell sections prior to electron imaging and is termed correlative light and electron microscopy (CLEM)^14^.

Here we present a modified strategy that combines correlative light microscopy and cryo-ET to locate regions of interest (ROI) in re-vitrified cell sections with subsequent determination of 3D structure by subtomogram averaging. We demonstrate the feasibility of this technique by determining structures of the herpes simplex virus type 1 (HSV-1) capsid, imaged within the nucleus of intact infected cells.

HSV-1 is an important human pathogen that causes cold sores. Infection with herpesviruses leads to life-long infection owing to their capacity to enter a latent state. More serious conditions caused by HSV-1 include encephalitis which may be fatal, and keratitis an eye condition that can lead to loss of sight ^15^. Herpesviruses have large double-stranded DNA genomes. Virion morphogenesis begins in the nucleus of infected cells with assembly of a T=16 icosahedral procapsid. Viral genomic DNA is pumped into the procapsid through a portal assembly located at a unique five-fold vertex^16^. The resulting mature nucleocapsid then buds through the nuclear envelope via an envelopment/de-envelopment process. In the cytoplasm, nucleocapsids become surrounded by a dense proteinaceous layer known as the tegument, before budding into plasma membrane derived endocytic membranes; giving rise to mature enveloped virions^17, 18^. The nucleocapsid structure has been extensively studied and was recently characterised at high resolution, leading to an atomistic model of this large (1250Å diameter) and complex assembly^19^. Of particular relevance to this study is the definition of the structure and composition of an assembly variously known as the capsid-vertex specific component (CVSC) or capsid associated tegument complex (CATC). Comprised of three proteins (pUL17, pUL25 and pUL36), this assembly forms a pentaskelion shaped structure that sits over the five-fold symmetry axes. Each arm of density is composed of a single copy of pUL17, and two copies of pUL25 and pUL36. Furthermore pUL36, described as an inner tegument protein, is understood to be the primary bridge between the capsid and tegument layers. The precise site of pUL36 addition to the capsids, in the nucleus or in the cytoplasm, has been the subject of much debate, with the balance of evidence suggesting that it binds capsids in the cytoplasm ^20, 21^. This is at odds with structural data that show two copies of pUL36 intimately associated with pUL17 and pUL25 to form a 5-helix bundle in the CATC^19^. pUL17 and pUL25 are definitively added in the nucleus. Our data reveal the presence of the CATC pentaskelion on HSV nucleocapsids in the nucleus, suggesting that capsids bind the tegument protein pUL36 (VP1/2) prior to nuclear egress.

## Results

### Correlative imaging of re-vitrified sections show<u>s</u> well-preserved ultrastructure

To image viral infectious processes within the cell, we used the vitreous frozen section (VFS) method. Virus infected cells were aldehyde fixed prior to being scraped and pelleted. The cell pellet was embedded in gelatin and cut into small cubes that were infused with sucrose. Embedded cells were then frozen in liquid nitrogen and ∼200 nm thick sections were cut at low temperature (−120°C). Frozen sections were collected on holey carbon film TEM grids and then washed to both warm the sections and remove the sucrose. Sections were then imaged at room temperature in the confocal microscope to identify regions of interest (ROIs), prior to being re-frozen by plunging into liquid ethane.

#### Correlative imaging of fluorescently tagged HSV-1 capsids in infected cells

Fluorescence light microscopy (FLM) was used to identify cells containing fluorescently tagged (RFP) wild type HSV-1 capsids, which were located in both the nucleus and cytoplasm. Sections were initially imaged in the laser-scanning confocal fluorescence microscope using tile mode to locate all sections on the grids (Supplementary Figure S1). Specific sections were chosen for further imaging in differential interference contrast (DiC) and fluorescence modes to obtain precise positions for HSV-1 capsids within cells (Supplementary Figure S1a). Sections were then re-vitrified and imaged by cryo-EM and cryo-ET. Initially the entire grid was imaged at low-magnification to detect the ROIs previously identified by light microscopy. We carried out image registration by overlaying the FLM and cryo-EM images to determine precise locations for subsequent cryo-ET (Figs 1a-b, Supplementary Figures S1b-d). Cryo-EM of the selected ROIs confirmed the presence of many HSV-1 capsids in the nucleus (Fig. 1d, 1f; Supplementary Fig. S1e) as well as in the cytoplasm (Fig. 1c, 1e; supplementary fig. S1e) of infected cells. Areas for cryo-ET were selected based on the presence of many capsids and minimal frost contamination and tomograms were recorded of capsids within the cytoplasm and the nucleus (Supplemental video 4). Our images therefore not only indicate good preservation of ultrastructure but also demonstrate the feasibility of using CLEM on re-vitrified sections to identify macromolecules of interest within the dense complex environment of the cell.

**Figure 1.**
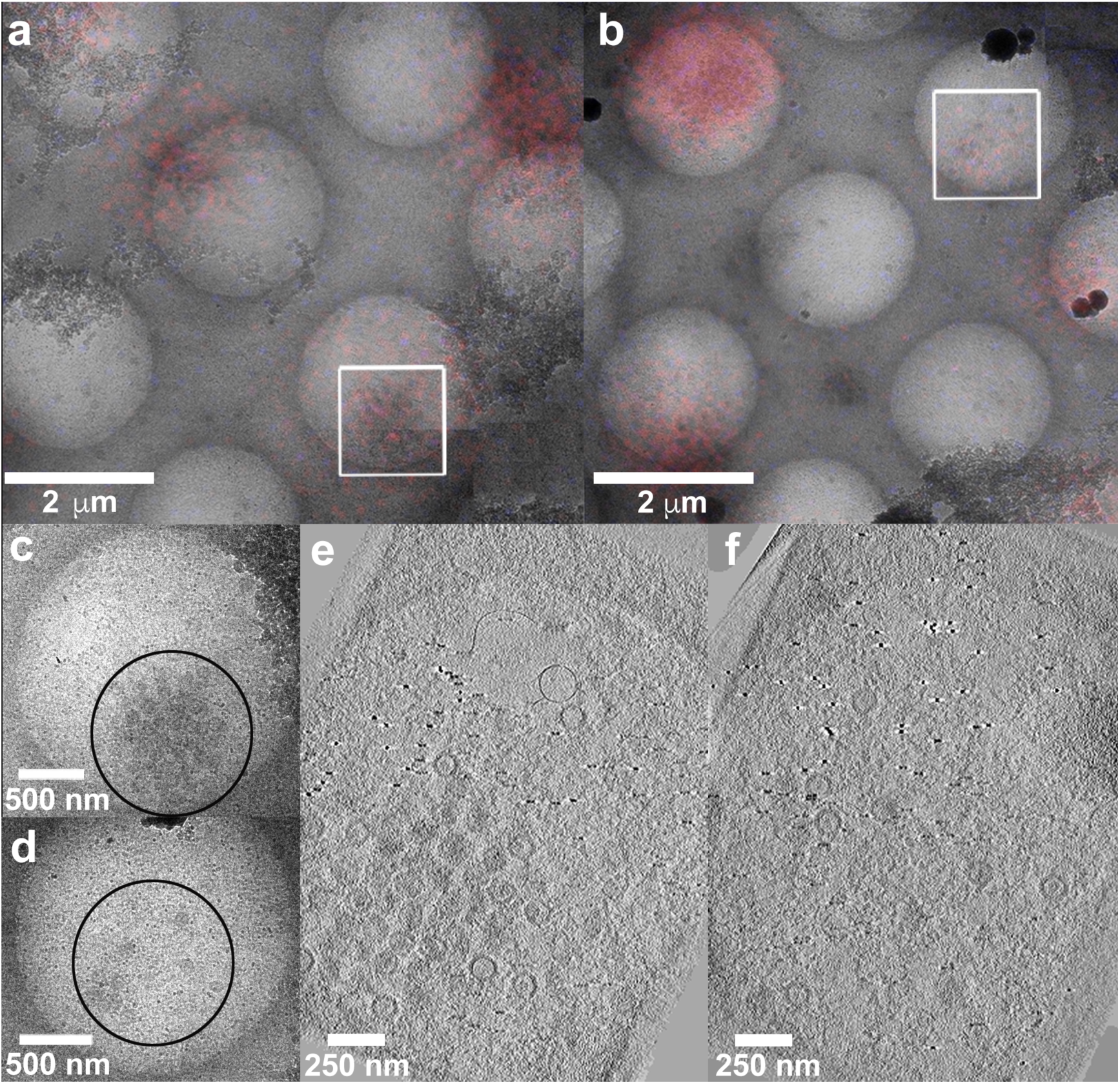
Correlative imaging of HSV virions. **(a) and (b)** Medium magnification light microscopy and EM images overlaid to find suitable ROI (white squares) with HSV capsids inside the cell. Wild type HSV-RFP tagged virions are shown (red). **(c)** High magnification EM image of the area of interest (white square in a) showing HSV capsids within the cytoplasm. **(d)** High magnification EM image of the area of interest (white square in b) denoting HSV capsids in the nucleus. **(e) and (f)** Z-slices through tomograms of cytoplasmic and nuclear capsids of areas denoted in c and d. Appropriate scale bars are shown.

### Cellular and capsid 3D ultrastructure revealed by cryo-ET of re-vitrified sections

The inner tegument protein pUL36 of HSV1 plays a vital role in virion assembly, however the site of its recruitment to the capsid, if carried out within the nucleus or outwith is unclear and has been a topic of much debate. To confirm this we imaged refrozen vitreous sections of cells infected with the mutant FRΔUL37 - capsids that lack the pUL37 tegument protein, but have pUL36. Cryo-EM images showed excellent preservation of cell ultrastructure combined with minimal cutting artefacts. At low magnification the nuclear membrane was readily seen, clearly demarcating the nucleus from the cytoplasm (Figure 2a-b).

**Figure 2.**
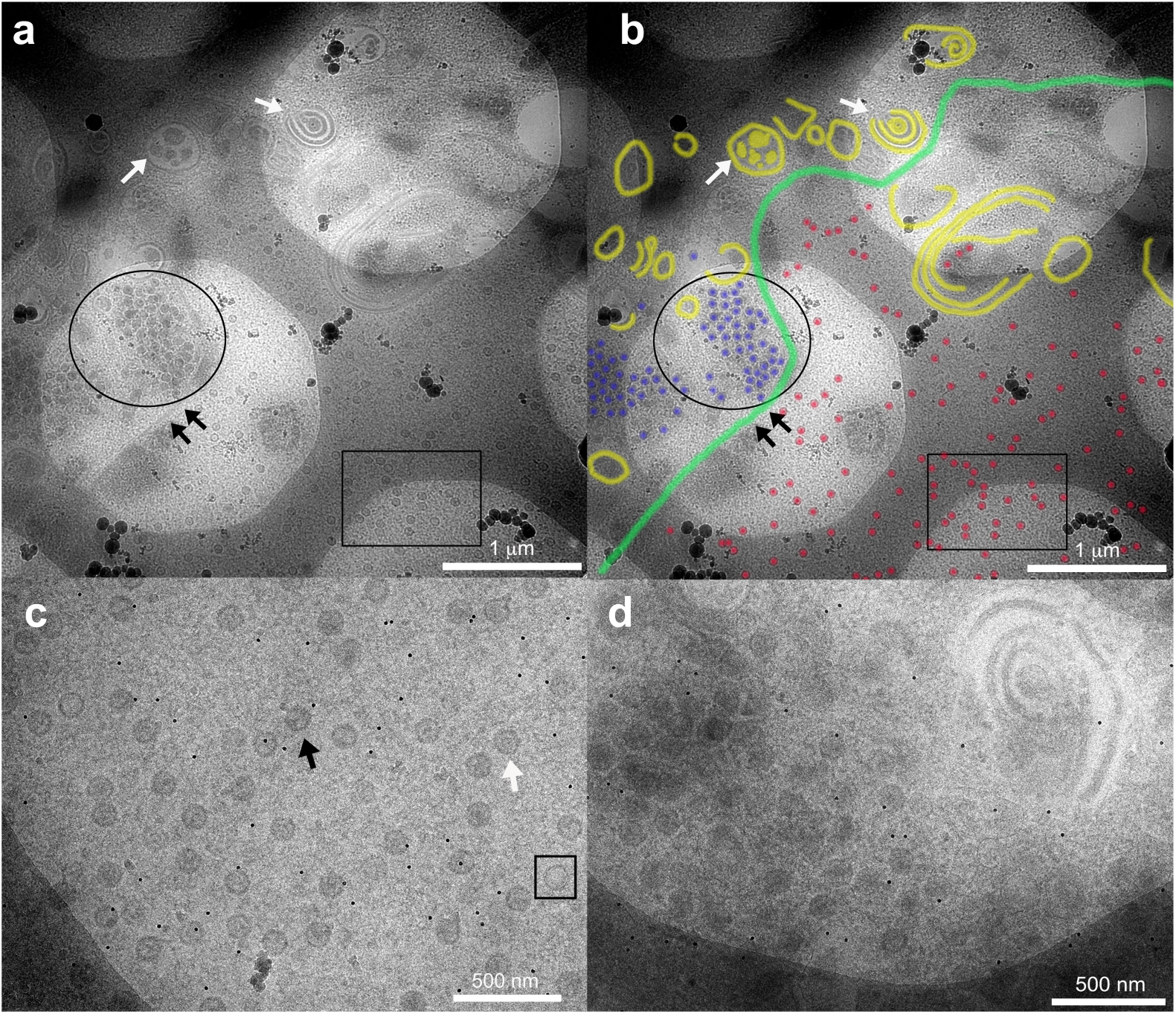
EM imaging of refrozen cell sections. **(a)** Low magnification diffraction imaging of a cell section by EM. Frozen sections show well-preserved structure and contrast. While an abundance of HSV capsids within the nucleus (black rectangle) and cytoplasm (black circle) are seen separated by the nuclear membrane (black arrows), other membranous and vacuolar structures (white arrows) can also be observed within the dense cytoplasm. **(b)** For easy visualisation, segmentation of the image in (a) using Photoshop is shown annotating the various structures namely, nuclear membrane (green), other membranous and vacuolar structures (yellow), nuclear capsids (red) and cytoplasmic capsids (blue). High magnification EM images of **(c)** nuclear and **(d)** cytoplasmic capsids clearly denote uniform dispersal of capsids within the nucleus, and a greater tendency to cluster up within the densely packed cytoplasm. One can also clearly observe the three different types of capsids (**in c**) namely, A-type (empty capsids, black square), B-type (scaffold-containing capsids, white arrow) and C-type (DNA-containing capsids, black arrow) within the nucleus. Appropriate scale bars are shown for the different images.

Several membranous and vacuolar structures were also visible within the cytoplasm. Cellular organelles such as mitochondria were observed and appeared to be well preserved (data not shown). We saw an abundance of uniformly distributed capsids within the nucleus, while in the cytoplasm the mutant capsids were found to be concentrated in dense aggregates as a result of the lack of pUL37 (Figure 2c-d). At higher magnification, the different types of capsids within the nucleus were easily discerned; empty A-capsids, scaffold-protein containing B-capsids and DNA filled C-capsids (Figs. 2, 3). We performed cryo-ET on capsid-rich areas within the nucleus and cytoplasm to generate 3D structure data. As expected, cryo-ET of capsids within the nucleus revealed an abundance of B- and C-capsids (Figure 3a-b, Supplemental video 5). Capsids in the cytoplasm were closely packed and were primarily C-capsids (Figure 3c-d).

**Figure 3.**
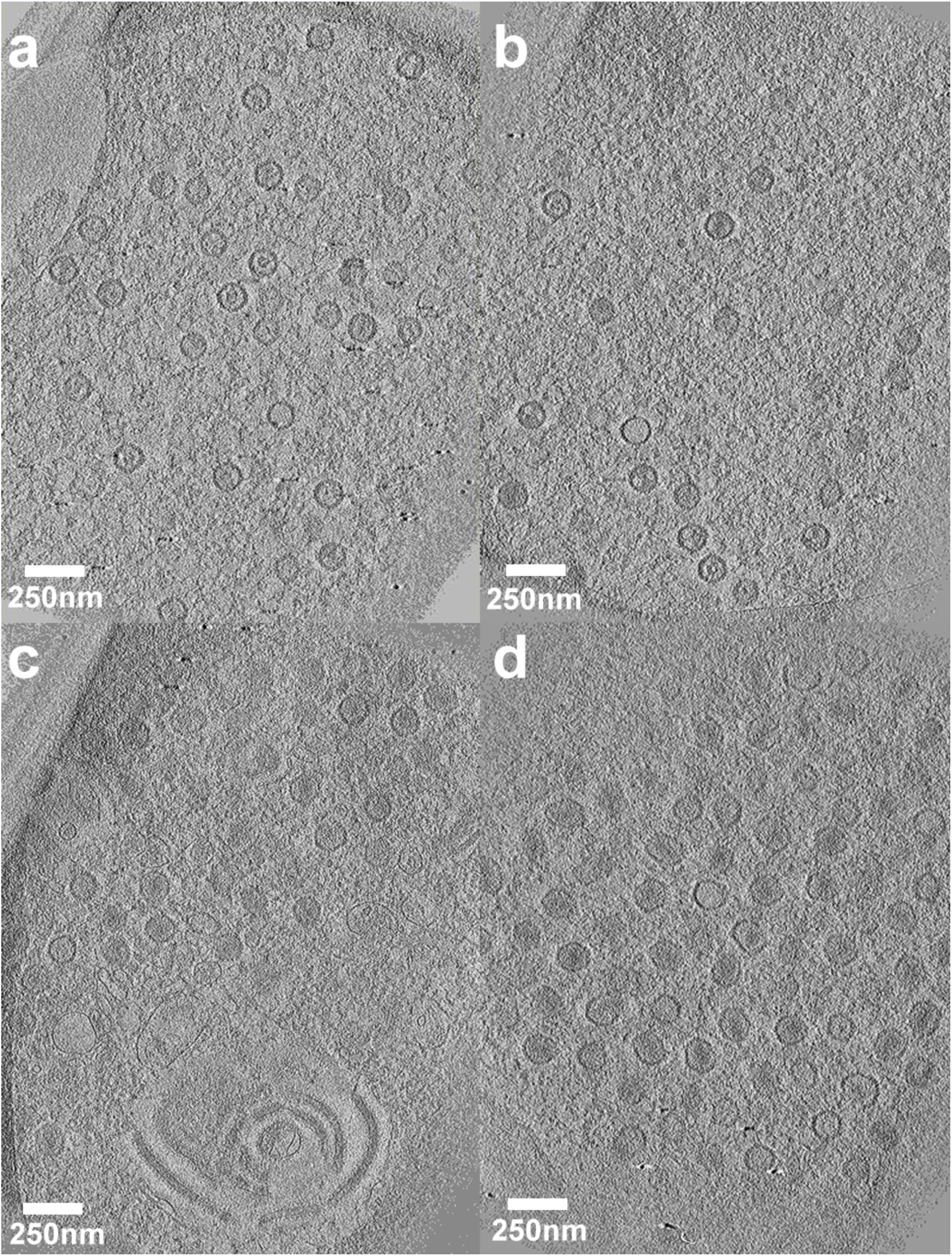
Electron tomography of nuclear and cytoplasmic HSV capsids. Tomograms were collected of the UL37-null mutant (FRΔUL37) capsids from within the nucleus and cytoplasm. While **(a) and (b)** represents a slice along the z-axis of the tomograms that show well dispersed areas of capsids within the nucleus, **(c) and (d)** are slices through tomograms showing capsids in the tightly-packed dense environment of the cytoplasm along with other membrane structures. Scale bar = 250 nm.

### 3D structure of the intranuclear HSV capsids by subtomogram averaging

To establish whether tomograms of refrozen cell section material are sufficient to determine 3D structures by sub-tomogram averaging, we sought to calculate *in-situ* reconstructions of HSV-1 capsids.

The inner tegument proteins pUL36 and pUL37 of HSV-1 play a critical role in virion assembly^22-24^. There has been uncertainty in the HSV field of how pUL36 and pUL37 are recruited to the capsid, if this happens within the nucleus or after nuclear egress^20, 21^. To shed light on this question, we carried out cryo-ET on the mutant lacking pUL37 (FRΔUL37). As capsids within the nucleus were discrete and well separated (Figure 3a-b), we performed subtomogram averaging to determine the 3D structures of the 3 classes of capsid present, i.e. A, B and C-capsids. In total 97 A-capsids 526 B-capsids and 155 C-capsids were selected and then processed by sub-tomogram averaging. Following several rounds of 3D classification and refinement along with imposition of icosahedral symmetry, reconstructions were calculated to a resolution of 5.7 nm (A-capsids), 5.6 nm (B-capsids) and 6.4 nm (C-capsids) for each class as validated by the Fourier shell correlation (FSC) assessments (Supplementary Fig. S2). Cross-sections of the structures obtained for each of the capsid classes showed clear differences in density between them; with A capsids being empty, while the B and C capsids harboured density within, corresponding to scaffold protein and DNA, respectively (Figure 4). Inspection of the capsid reconstructions revealed differences in CATC density observed at the 5-fold vertices between the three subclasses; the pentaskelion structure is clearly visible in the C-capsids (Figure 4, Supplemental video 6).

**Figure 4.**
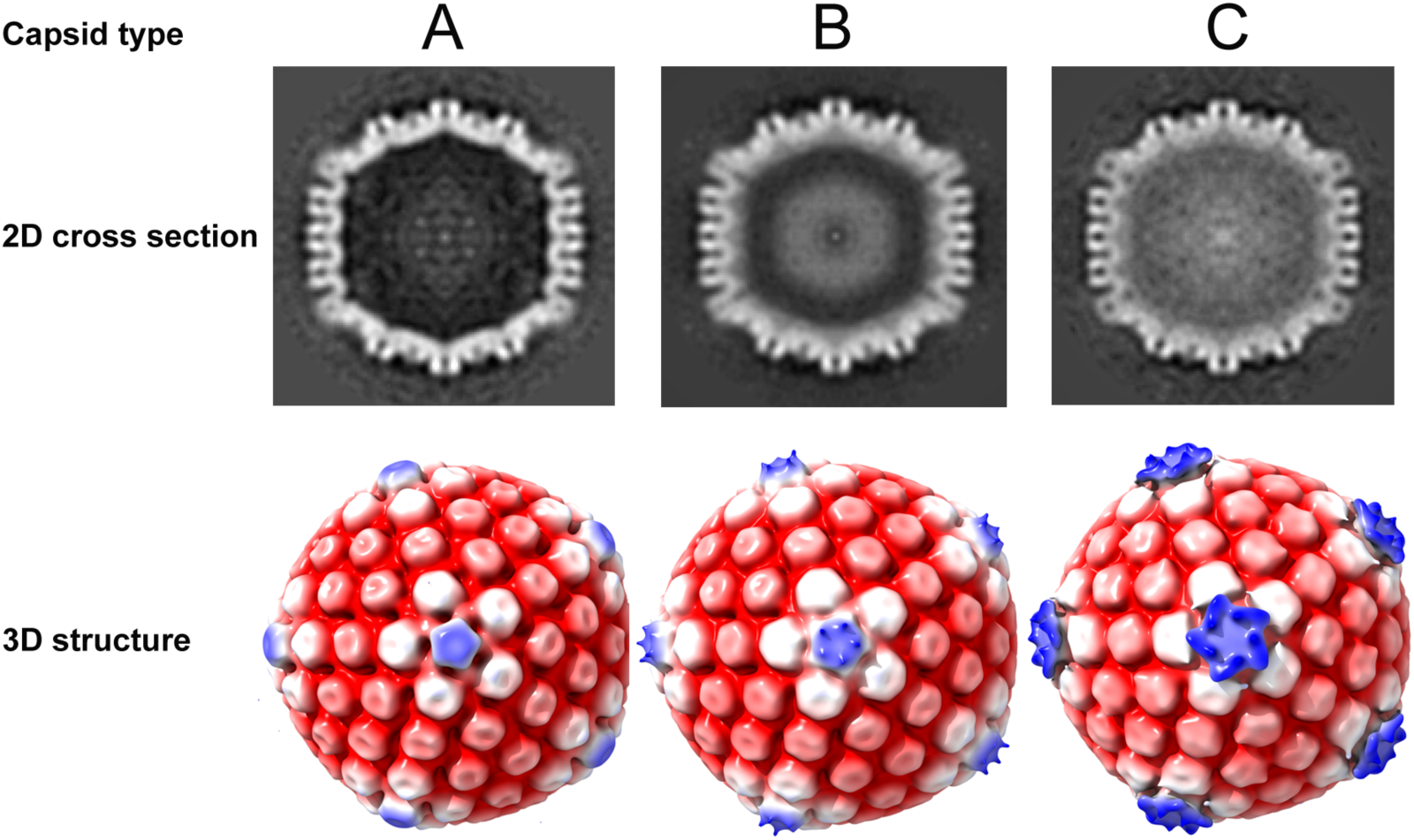
Determination of the 3D HSV capsid structures from within the nucleus. 3D structures of the three types of nuclear capsids, A-, B- and C-type were determined by subtomogram averaging. 2D cross-section images clearly indicate density differences in the interior of the capsid between the three types. Density relating to scaffolding proteins and DNA is seen inside the capsids within the B-type and C-type, respectively, while none is present inside the empty capsid shell of the A-type. The 3D structures obtained have been radially coloured at low thresholds of density and are shown along the 5-fold axis of icosahedral symmetry. Although additional pentaskelion density is observed above the pentons in both B and C-type capsids, it is more pronounced in the latter.

In recent high-resolution single particle reconstructions of capsids within purified HSV-1 virions, the pentaskelion structure can be clearly seen above the penton vertices and has been attributed to the proteins pUL17, pUL25 and pUL36^16, 19^. Although at low-resolution, the star-shaped density observed in our C-capsid structure is strongly reminiscent of the penton density in the high-resolution structures determined from purified HSV-1 virions (Figure 5).

**Figure 5.**
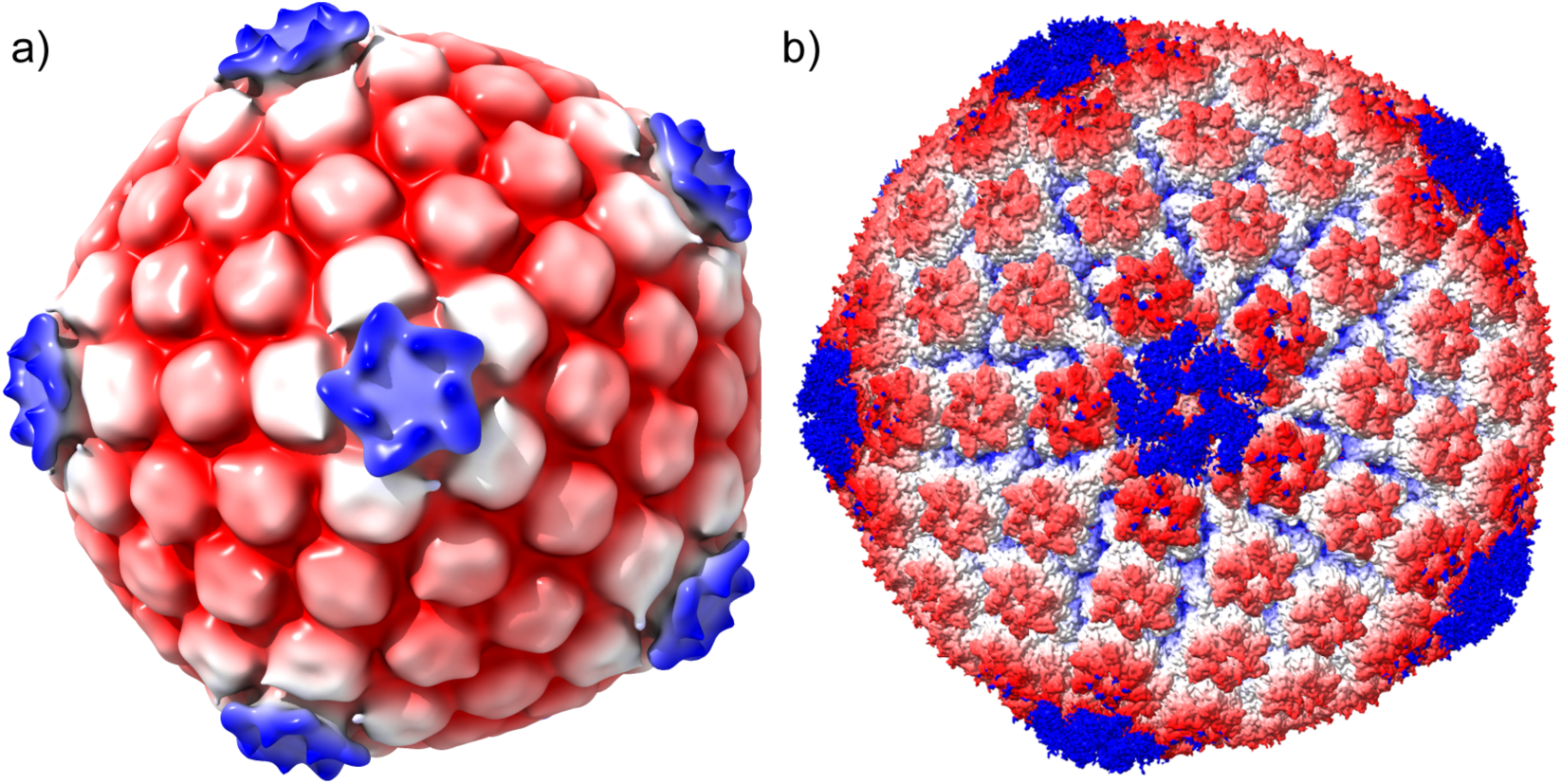
Comparison of *in situ* intranuclear capsid structure with the high-resolution structure of purified HSV-1 capsid. **(a)** The *in situ* 3D structure of C-capsid determined by subtomogram averaging at 6.4 nm shows clear pentaskelion electron density above the pentons at low thresholds of density. This corresponds to the CATC, reported to be composed of pUL36, pUL17 and pUL25 proteins; and is comparable to the **(b)** high-resolution structure of purified capsids from within the nucleus16. Capsids are radially coloured from the centre.

The CATC density was hardly visible in the B-capsids. We attribute this to low occupancy of CATC in this class of capsids, resulting in its signal being lost during particle and symmetric averaging. The CATC density is completely absent in the A-capsids.

Focussed classification has recently emerged as a means to obtain structural data pertaining to low-occupancy, unique and asymmetric features that are otherwise lost owing to particle averaging upon imposition of higher order symmetry during computational processing^25^. We successfully implemented this method and determined the structure of the unique portal vertex in purified HSV-1 virions^16^. To assess if we might similarly reveal asymmetric features such as the HSV-1 portal or additional density at the penton vertices due to low-occupancy, we carried out focussed classification on a single 5-fold vertex of the 3D *in situ* structures of the capsids (A, B and C-type) obtained from subtomogram averaging. To our knowledge this is the first attempt to implement this method on sub tomograms. Focussed classification allows for expansion of symmetry, where each virus particle that would normally have a single orientation defined relative to the asymmetric unit, will now encompass 60 symmetry-related orientations of the icosahedral capsid. Thus, each five-fold vertex may be individually masked and classified. Although a total of 10 classes were calculated, only 1 class was identified as containing a distinct star-shaped structure in B- and C-capsids, corresponding to CATC density (Figure 6, Supplementary Fig. S3).

**Figure 6.**
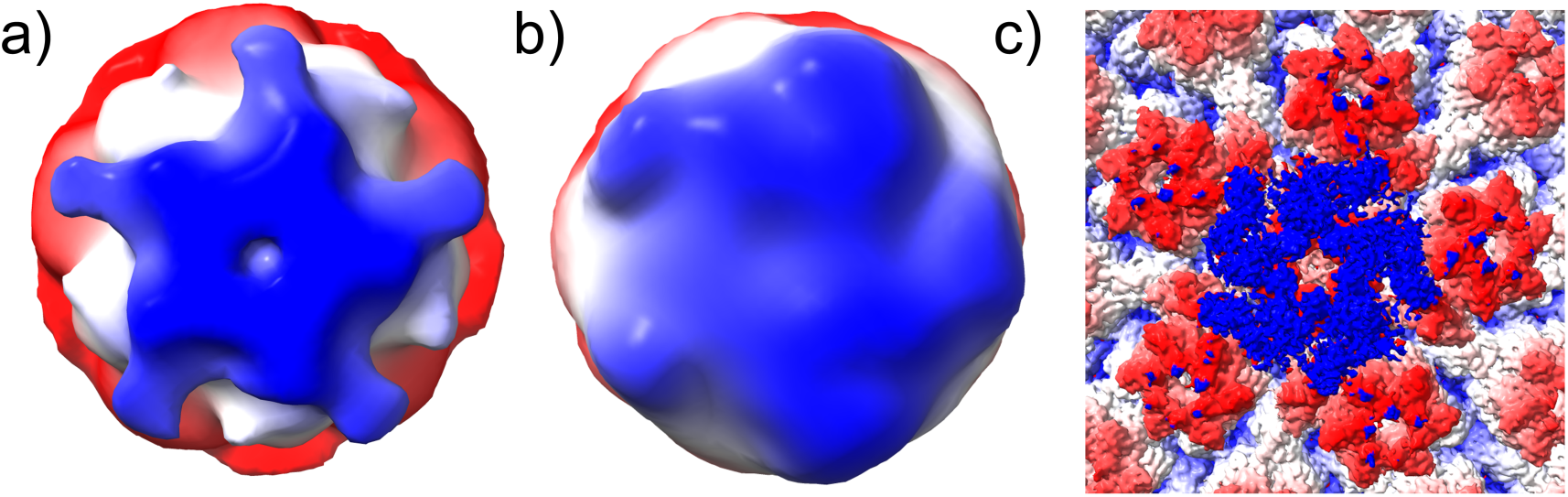
Asymmetric focussed classification of intranuclear capsids. The 3D structures of the most meaningful class obtained from focused classification of (**a)** B and (**b)** C-type capsids indicate distinct density above the 5-fold vertex resembling a pentaskelion. However, the density on the C-capsid is less clear owing to the reduced number of particles used during the focused classification step. **(c)** Comparison with a close up image of the penton density from the high-resolution structure of purified nuclear capsids, confirmed unequivocally the density to pertain to CATC that is present on matured capsids within the nucleus.

None was observed in the classes calculated from the A-capsids (data not shown). The pentaskelion density in B-capsids is more prominent than C-capsids; likely owing to far greater numbers of B-capsids (526) used during processing than of C-capsids (125). These data support our suggestion that low occupancy of CATC on B-capsids led to weaker density in icosahedrally averaged density maps. They are clearly visible upon asymmetric reconstruction (Figure 6), but not during symmetric reconstruction (Figure 4). Focussed classification of all three capsid subgroups (A, B and C) however did not reveal any structure pertaining to the portal. This is likely owing to low overall numbers of particles used for the calculations.

## Discussion

Cryo-EM is fast becoming a critical tool for structural biologists, allowing atomic-resolution structures of proteins and viruses to be calculated rapidly^2^. In this work, we have addressed the main bottleneck of obtaining 3D structures of macromolecular assemblies from within the native context of the cell under cryo conditions. By adapting the method of classical Tokuyasu sectioning coupled with CLEM and subtomogram averaging, we offer an easy and attractive alternative for imaging and 3D structure determination of proteins/viruses from within cells^9^. The Tokuyasu method has been previously used to produce sections that were subsequently vitrified for cryo-EM imaging^10^. However, the use of vitrified frozen sections (VFS) has thus far been limited to imaging tissue sections by cryo-EM and cryo-ET^10-12^.

We believe our method offer a less technically challenging and relatively inexpensive alternative for the majority of cryo-EM users to carry out 3D structural characterisation of macromolecular assemblies in cellular and tissue samples. Further, the option of using CLEM on these sections offers the great advantage of precisely locating sub-cellular structures of interest, broadening the applicability of this method to the study of low-frequency events. 3D structure determination of macromolecular assemblies from re-frozen sections via subtomogram averaging provides a starting point to more routine high-resolution structure determination of proteins from within the cell.

The use of chemical fixative may cause some structural artefacts, possibly contributing to the low resolution of capsid structures in our study (5-6 nm), in comparison to resolutions obtained from subtomogram averaging of proteins from unfixed cryo-ET of for example purified virions (0.8-2 nm). In the absence of comparable data from FIB-SEM prepared nuclear capsids, it is not possible to directly compare achievable results and thereby unambiguously assign reasons for the low resolution achieved here.

In this study we chose the highly symmetric and well characterized HSV capsid as a model to test the feasibility of our method to determine the *in situ* 3D structures of intracellular capsids by subtomogram averaging. Employing a correlative approach using CLEM, allowed us to spatially locate capsids within both the nucleus and cytoplasm. Well-preserved fluorescence signal as well as ultrastructure within the sections observed was consistent with previously published work describing correlative imaging of refrozen Tokuyasu sections ^13^.

A complex variable proteinaceous tegument layer, comprising more than 20 proteins surrounds the HSV capsid. Of these, pUL36 and pUL37 are classified as ‘inner-tegument’ proteins and are tightly associated with the capsid. The precise location of pUL37 has not been determined. pUL36 is intimately associated with pUL25 and pUL17 to form the CATC at the five-fold vertices^26-28^. With the pUL37-null mutant used in this work, accumulation of C-capsids in dense clusters was observed in the cytoplasm of infected cells, confirming that the tegument protein pUL37 is not important for capsid assembly, DNA packaging or nuclear egress, but plays a vital role in tegument addition within the cytoplasm and envelopment. Despite extensive studies on HSV-1 capsid morphogenesis, controversy persists concerning the cellular compartment in which the tegument protein pUL36 is added to the nascent virus capsid. The CATC density observed at capsid five-fold vertices in cryo-EM studies of mature virions, includes a five-helix bundle that is the principal site of interaction for two copies each of pUL36 and pUL25 along with a single copy of pUL17 ^20, 26, 28-31^. The intranuclear capsid structures obtained from subtomogram averaging here clearly indicate variable pentaskelion CATC density over the pentons between the three subgroups of capsids; B-capsids having weak density compared to C-capsids, likely a consequence of low occupancy of CATC in B-capsids. Upon performing focussed classification for both B- and C-capsids we revealed the presence of CATC density at the penton vertex, comparable to previously reported high-resolution structures of capsids within purified HSV-1 virions^16, 19^. Although the presence of putative pUL36 in the pentaskelion density of these low-resolution capsid structures cannot be confirmed, our data strongly argue for a fully assembled CATC prior to nuclear egress. We cannot rule out the proposed incorporation of N-terminally truncated forms of pUL36 however ^32-34^.

In summary, we have demonstrated a relatively inexpensive and technically straight-forward method for determining *in situ* 3D structures of macromolecular assemblies using CLEM and subtomogram averaging. Despite the wide applicability of this method, it could be improved and optimised for the future; automatic data collection of large datasets using 300 KV microscopes with high-speed direct electron detectors would be an obvious starting step. This may enable determination of high-resolution structures from re-vitrified sections of cells and tissues by subtomogram averaging techniques. Finally, we validated this method by determining the first structures of icosahedral capsids directly determined from within the nucleus. Our method opens the possibility of determining and characterising specific complexes and their interactions at high-resolution within the functional context of the cell or tissue, providing snapshots of important and dynamic events in biology.

## Author Contributions

The CVR has adopted the CRediT taxonomy (https://casrai.org/credit/). Author’s contribution is as follows. SV: Conceptualization, Project administration, Investigation, Methodology, Formal analysis, Visualization, Writing - original draft, review & editing. FJR: Conceptualization, Investigation, Methodology, Resources; MM: Investigation, Resources; CL: Investigation, Resources, Methodology; DB: Conceptualization, Investigation, Methodology, Formal analysis, Funding acquisition, Writing – review & editing.

## Materials and methods

### Cells and viruses

Baby hamster kidney cells (BHK-21) were cultured as described previously^23, 35^. Briefly, cells were grown at 37°C in Glasgow minimum essential medium (GMEM; Invitrogen) supplemented with 10% newborn calf serum (NCS; Invitrogen) and tryptose phosphate broth (TPD; Invitrogen). BHK-21 cells were infected using wild-type HSV-1 (strain 17^+^) pUL35 RFP or UL37-null (FRΔUL37) virus at 10 PFU per cell and incubated for fixed times post infection.

### Preparation of HSV infected cells for sectioning

Petri dishes (60/100 mm) dishes were cultured with BHK-21 cells and grown to confluency prior to infection with wild type pUL35RFP1D1 (pUL35 tagged with the RFP fluorophore) or FRΔUL37 strains of HSV at 10 PFU/cell for 12h and 9h post infection, respectively. Cells were then washed with PBS and fixed with 2% formaldehyde and 1% glutaraldehyde in PBS for 5h at room temperature or at 4°C overnight. The following day, cells were washed with PBS, scraped, pelleted and resuspended in 12% porcine gelatin (Sigma) and cooled on ice. Cooled pellets were cut into 1 mm cubes and infiltrated with 2.3 M sucrose at 4°C for 2-3 days. Well-infiltrated blocks were attached to Tokuyasu stubs (Leica Microsystems, Germany), frozen in liquid nitrogen and stored in cryotubes in liquid nitrogen dewars until future use.

### Tokuyasu sectioning

Sucrose-infiltrated frozen cell blocks were transferred to the cryo ultramicrotome (Ultracut EM FC6, Leica Microsystems). 200-220 nm thick sections were cut with a cryo-immuno diamond knife (Diatome) and were collected using a “Perfect loop” (Diatome) with 2.3 M sucrose^9^ and transferred to holy carbon finder grids (R3.5/1 Copper NH2 - Quantifoil, Jena, Germany). Sections were stored at 4 °C until further processing.

### Correlative confocal light microscopy

Cut sections were washed with PBS to remove all traces of sucrose. Prior to imaging, DAPI (4′,6-diamidino-2-phenylindole dihydrochloride, Sigma, 1:500 dilution) was applied for 30 min followed by washes in PBS (3 × 10 sec). Sections on the grid were then transferred to glass-bottom dishes (Mattek Corp, USA) on ice with sections facing down submerged in minimal buffer (∼800 μl) and imaged. Confocal laser scanning microscopy was performed on the LSM 880 microscope (Zeiss, Germany) with a 63x/1.4 NA oil immersion objective. Tiled images (20×20 tile) of the entire grid as well as single images and image stacks (2048 × 2048 × 5) were acquired with a pinhole close to 1 Airy unit. DAPI (405 nm excitation) and RFP (530 nm excitation) signals were acquired with GaAsP detectors (Zeiss, Germany). Image processing and deconvolution was carried out with the Zeiss licenced software, ZEN Black.

### Cryo EM/ET and tomogram reconstruction

Sections imaged by light micoscopy were re-vitrified and prepared for cryogenic transmission electron microscopy using an FEI Vitrobot Mk IV. Briefly, 6-8 μl of 15 nm colloidal gold (1:3 dilution, BBI, UK) was applied to the finder grids with sections and blotted for 5 seconds before plunging into liquid ethane^36^. Vitrified samples were viewed at low-temperature (around 98 K) and under low electron dose conditions using a JEOL 2200 FS cryo-microscope operated at 200 KV with zero-loss energy filtered imaging using a slit-width of 30 eV. Samples were held in a Gatan 914 high-tilt cryo-stage. CEM images were typically recorded at zero tilt at low magnifications of 100X-200X as well as medium magnifications of 6000X-8000X using the Gatan Ultrascan 4k × 4k CCD camera to determine the same ROIs and enable superposition with light microscopy images. Tilt series were then recorded using the SerialEM software package^37^ at 10,000 × magnification on a Direct-Electron DE20 DDD camera at a sample rate of 5.98 Å/pixel. Images were acquired at two-degree increments with a bi-directional tilt scheme ranging from −20° to +60° for the first tilt set followed by −20° to −60° for the second tilt set. The target defocus was set between 4-10 μm and the electron dose ranged from 50-100 e/Å^2^ per tilt-series. Tomograms were calculated and visualized using the IMOD software package^38^. Reconstruction was performed using weighted back projection followed by CTF correction available as part of the IMOD package.

### Registration of LM and EM images

Alignment of light and electron microscopy images was done with Adobe Photoshop CS5 software. Firstly, the LM image was aligned based on the markings of the finder grid, manually inserted landmarks such as ice contamination and the nuclear DAPI signal and their corresponding signals on the cryo-EM image. After rough alignment, a fine alignment was performed by registering the pattern of holes within the grid square and/or the pattern of cells and their corresponding signals on the cryo-EM image. For data presentation, the light and electron microscopy images were merged with the merge channels option in Adobe photoshop CS5.

### Computational subtomogram image processing

Micrograph movies containing 20 frames were processed to calculate a 3D reconstruction using Relion-3.1^39^. Initially movies were corrected for specimen movement and radiation damage using Direct-Electron open source software scripts (DE_combine_references.py and DE_process_frames.py). Processed images were stacked using in-house scripts and tomograms generated with defocus estimation (Ctfplotter) using IMOD^38^. An initial set of 778 particles from 33 tomograms of nuclear capsids was manually picked and extracted using the EMAN program e2boxer.py^40^. Extracted particles were grouped separately based on them being A, B or C-type capsids prior to classification. 3D classification was performed in Relion-3.1 imposing icosahedral symmetry. For the 3D classification, a low resolution starting model of the HSV capsid, obtained after processing and refining ∼100 manually extracted particles from our tomogram dataset via EMAN2 was used^39, 40^. The subvolumes and the starting model were 2x binned to speed up processing. A subset of particles from each of the three subgroups (A, B, and C-type capsids) was selected from the 3D class that yielded the highest resolution and subjected to further refinement. Resolution assessment was done with Relion, using the postprocessing task to mask the density maps and calculate the ‘gold-standard’ Fourier shell correlation. The final 3D reconstructions of A, B and C-type capsids were visualized in UCSF Chimera, ChimeraX and IMOD^38, 41, 42^ Movies were created using ChimeraX^42^.

Following 3D reconstruction, focused classification was performed on each of the capsid subgroups (A, B and C-type capsids) using Relion-3.1 to identify unique or additional density on the 5-fold vertex^16^. A cylindrical mask was created in SPIDER covering a single 5-fold vertex^43^. Symmetry expansion of the dataset was performed in Relion by creating a new metadata file that comprised of information pertaining to 60 orientations for each virus particle. The dataset was 2x binned to speed up calculations. The subvolumes were subjected to 3D classification with a T value of 5, to reconstruct a single 5-fold vertex, without refining orientations and origins. A total of 10 classes were calculated with one of them identified to have apparent pentaskelion density over the 5-fold axis, corresponding to CATC, in both B-capsids and C-capsids. All visualization was carried out using ChimeraX^42^.

## References

1. Mollenhauer, H.H. Artifacts caused by dehydration and epoxy embedding in transmission electron microscopy. Microsc Res Tech 26, 496–512 (1993).

2. Chang, J., Liu, X., Rochat, R.H., Baker, M.L. & Chiu, W. Reconstructing virus structures from nanometer to near-atomic resolutions with cryo-electron microscopy and tomography. Adv Exp Med Biol 726, 49–90 (2012).

3. Loney, C., Mottet-Osman, G., Roux, L. & Bhella, D. Paramyxovirus ultrastructure and genome packaging: cryo-electron tomography of sendai virus. J Virol 83, 8191–8197 (2009).

4. Vijayakrishnan, S. et al. Cryotomography of budding influenza A virus reveals filaments with diverse morphologies that mostly do not bear a genome at their distal end. PLoS Pathog 9, e1003413 (2013).

5. Oikonomou, C.M. & Jensen, G.J. Cellular Electron Cryotomography: Toward Structural Biology In Situ. Annu Rev Biochem 86, 873–896 (2017).

6. Al-Amoudi, A., Norlen, L.P. & Dubochet, J. Cryo-electron microscopy of vitreous sections of native biological cells and tissues. J Struct Biol 148, 131–135 (2004).

7. Al-Amoudi, A., Studer, D. & Dubochet, J. Cutting artefacts and cutting process in vitreous sections for cryo-electron microscopy. J Struct Biol 150, 109–121 (2005).

8. Marko, M., Hsieh, C., Schalek, R., Frank, J. & Mannella, C. Focused-ion-beam thinning of frozen-hydrated biological specimens for cryo-electron microscopy. Nat Methods 4, 215–217 (2007).

9. Tokuyasu, K.T. A technique for ultracryotomy of cell suspensions and tissues. J Cell Biol 57, 551–565 (1973).

10. Sabanay, I., Arad, T., Weiner, S. & Geiger, B. Study of vitrified, unstained frozen tissue sections by cryoimmunoelectron microscopy. J Cell Sci 100 (Pt 1), 227–236 (1991).

11. Bokstad, M., Sabanay, H., Dahan, I., Geiger, B. & Medalia, O. Reconstructing adhesion structures in tissues by cryo-electron tomography of vitrified frozen sections. J Struct Biol 178, 76–83 (2012).

12. Luther, P.K. & Morris, E.P. Cryoelectron microscopy of refrozen cryosections. J Struct Biol 142, 233–240 (2003).

13. Bos, E. et al. Vitrification of Tokuyasu-style immuno-labelled sections for correlative cryo light microscopy and cryo electron tomography. J Struct Biol 186, 273–282 (2014).

14. Bykov, Y.S., Cortese, M., Briggs, J.A. & Bartenschlager, R. Correlative light and electron microscopy methods for the study of virus-cell interactions. FEBS Lett 590, 1877–1895 (2016).

15. Whitley, R.J. in Fields’ Virology, Vol. 2. (eds. D.M. Knipe & P.M. Howley) 2461–2509 (Lippincott Williams & Wilkins, Philadelphia; 2001).

16. McElwee, M., Vijayakrishnan, S., Rixon, F. & Bhella, D. Structure of the herpes simplex virus portal-vertex. PLoS Biol 16, e2006191 (2018).

17. Hollinshead, M. et al. Endocytic tubules regulated by Rab GTPases 5 and 11 are used for envelopment of herpes simplex virus. EMBO J 31, 4204–4220 (2012).

18. Roizman, B., Knipe, D.M. & Whitley, R.J. in Fields Virology, Vol. 2, Edn. 6. (eds. D.M. Knipe et al.) 1824–1897 (Lippincott Williams & Wilkins, Philadelphia; 2013).

19. Dai, X. & Zhou, Z.H. Structure of the herpes simplex virus 1 capsid with associated tegument protein complexes. Science 360 (2018).

20. Bucks, M.A., O’Regan, K.J., Murphy, M.A., Wills, J.W. & Courtney, R.J. Herpes simplex virus type 1 tegument proteins VP1/2 and UL37 are associated with intranuclear capsids. Virology 361, 316–324 (2007).

21. Henaff, D., Remillard-Labrosse, G., Loret, S. & Lippe, R. Analysis of the early steps of herpes simplex virus 1 capsid tegumentation. J Virol 87, 4895–4906 (2013).

22. Kelly, B.J. et al. The interaction of the HSV-1 tegument proteins pUL36 and pUL37 is essential for secondary envelopment during viral egress. Virology 454-455, 67–77 (2014).

23. Roberts, A.P. et al. Differing roles of inner tegument proteins pUL36 and pUL37 during entry of herpes simplex virus type 1. J Virol 83, 105–116 (2009).

24. Desai, P.J. A null mutation in the UL36 gene of herpes simplex virus type 1 results in accumulation of unenveloped DNA-filled capsids in the cytoplasm of infected cells. J Virol 74, 11608–11618 (2000).

25. Conley, M.J. & Bhella, D. Asymmetric analysis reveals novel virus capsid features. Biophys Rev 11, 603–609 (2019).

26. Grunewald, K. et al. Three-dimensional structure of herpes simplex virus from cryo-electron tomography. Science 302, 1396–1398 (2003).

27. Mettenleiter, T.C. Herpesvirus assembly and egress. J Virol 76, 1537–1547 (2002).

28. Zhou, Z.H., Chen, D.H., Jakana, J., Rixon, F.J. & Chiu, W. Visualization of tegument-capsid interactions and DNA in intact herpes simplex virus type 1 virions. J Virol 73, 3210–3218 (1999).

29. Cardone, G. et al. The UL36 tegument protein of herpes simplex virus 1 has a composite binding site at the capsid vertices. J Virol 86, 4058–4064 (2012).

30. Cockrell, S.K., Huffman, J.B., Toropova, K., Conway, J.F. & Homa, F.L. Residues of the UL25 protein of herpes simplex virus that are required for its stable interaction with capsids. J Virol 85, 4875–4887 (2011).

31. Toropova, K., Huffman, J.B., Homa, F.L. & Conway, J.F. The herpes simplex virus 1 UL17 protein is the second constituent of the capsid vertex-specific component required for DNA packaging and retention. J Virol 85, 7513–7522 (2011).

32. Leelawong, M., Lee, J.I. & Smith, G.A. Nuclear egress of pseudorabies virus capsids is enhanced by a subspecies of the large tegument protein that is lost upon cytoplasmic maturation. J Virol 86, 6303–6314 (2012).

33. Schipke, J. et al. The C terminus of the large tegument protein pUL36 contains multiple capsid binding sites that function differently during assembly and cell entry of herpes simplex virus. J Virol 86, 3682–3700 (2012).

34. Wan, W. et al. Structure and assembly of the Ebola virus nucleocapsid. Nature 551, 394–397 (2017).

35. Fan, W.H. et al. The large tegument protein pUL36 is essential for formation of the capsid vertex-specific component at the capsid-tegument interface of herpes simplex virus 1. J Virol 89, 1502–1511 (2015).

36. Adrian, M., Dubochet, J., Lepault, J. & McDowall, A.W. Cryo-electron microscopy of viruses. Nature 308, 32–36 (1984).

37. Mastronarde, D.N. Automated electron microscope tomography using robust prediction of specimen movements. J Struct Biol 152, 36–51 (2005).

38. Kremer, J.R., Mastronarde, D.N. & McIntosh, J.R. Computer visualization of three-dimensional image data using IMOD. J Struct Biol 116, 71–76 (1996).

39. Scheres, S.H. RELION: implementation of a Bayesian approach to cryo-EM structure determination. J Struct Biol 180, 519–530 (2012).

40. Tang, G. et al. EMAN2: an extensible image processing suite for electron microscopy. J Struct Biol 157, 38–46 (2007).

41. Pettersen, E.F. et al. UCSF Chimera--a visualization system for exploratory research and analysis. J Comput Chem 25, 1605–1612 (2004).

42. Goddard, T.D. et al. UCSF ChimeraX: Meeting modern challenges in visualization and analysis. Protein Sci 27, 14–25 (2018).

43. Frank, J. et al. SPIDER and WEB: processing and visualization of images in 3D electron microscopy and related fields. J Struct Biol 116, 190–199 (1996).

