## Supplemental Figures for "*In situ* structure determination of virus capsids imaged within cell nuclei by correlative light and cryo-electron tomography"

**Supplemental data**

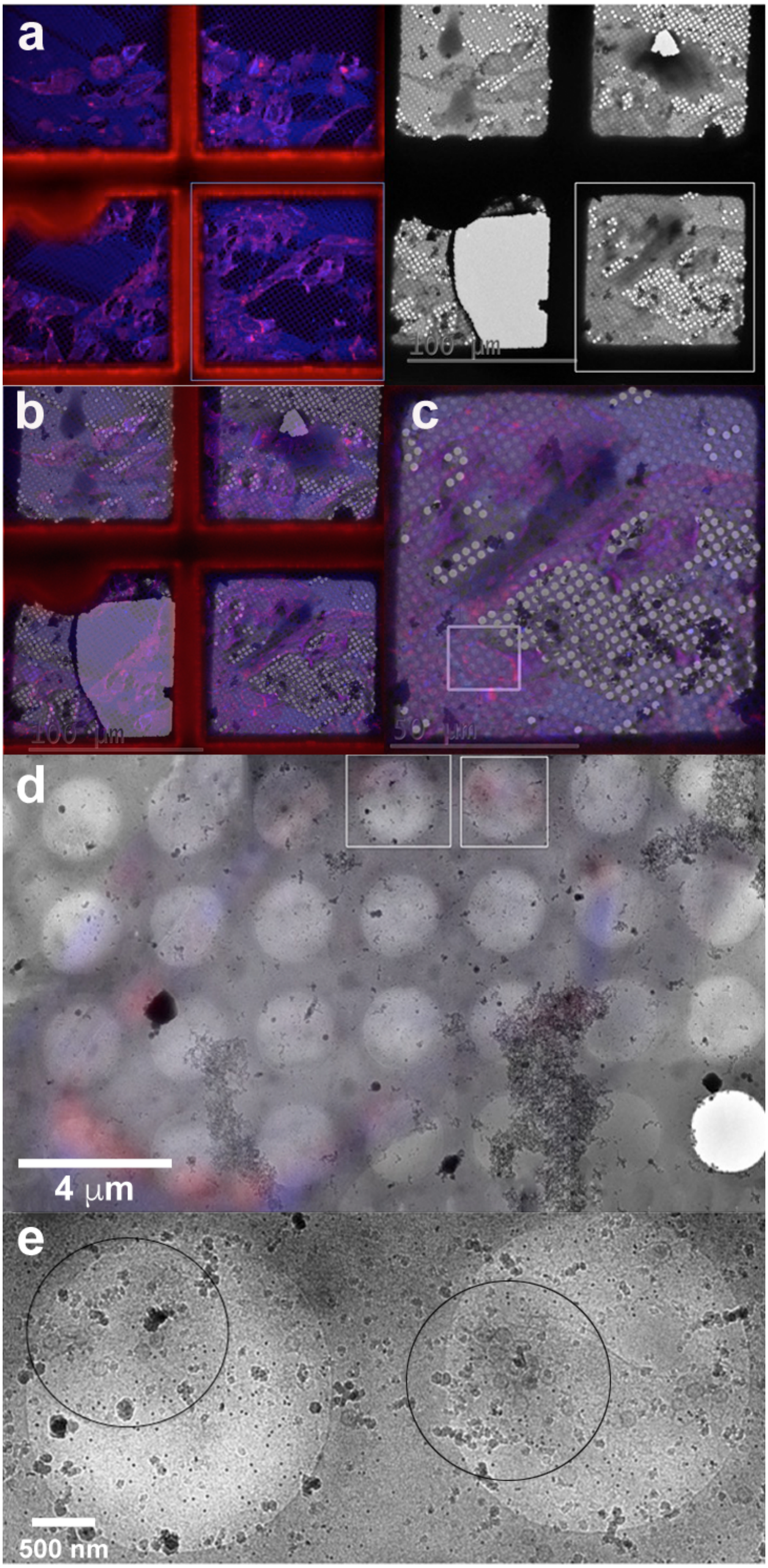
a
100 μm
b
100 μm
c
50 μm
d
4 μm
e
500 nm

### Figure S1. Correlative imaging of HSV capsids.

Overlay of LM and EM images at low magnifications to obtain areas of interest pertaining to wild type HSV capsids in cells, both within the nucleus and cytoplasm. **(a)** On the left is a low magnification confocal image of an area within the cell section depicting RFP tagged HSV (red) infected cells. Nuclei are stained with DAPI (blue). The right side is the equivalent area imaged by EM at low magnifications. **(b)** Image registration and overlay of both low-magnification LM and EM images shown in (a). **(c)** Close up of the LM-EM overlaid image of the interested grid square (white square of (a)) showing the chosen position of a cell in greater detail (white square) for further CET analysis **(d)** LM and EM image superposition at medium magnifications of the chosen cell depicted in the white square of (c). High-magnification **(e)** EM images of regions selected in (d) shown as white squares reveal HSV capsids present in both the nucleus and cytoplasm.

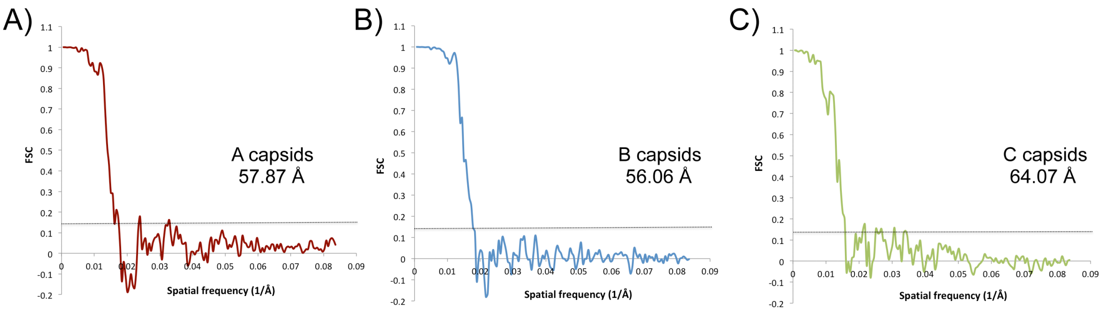


### Figure S2. Resolution assessment of HSV capsid structures.

Resolution assessment of 3D reconstructions of virus capsids was determined by the gold standard Fourier shell correlation (FSC) analysis, as part of the Relion-3.1 package. Graphs denoting the resolution of capsids calculated at the FSC 0.143 cutoff to be **(A)** 57.87 Å for A-capsids, **(B)** 56.06 Å for B-capsids and **(C)** 64.07 Å for C-capsids are shown.

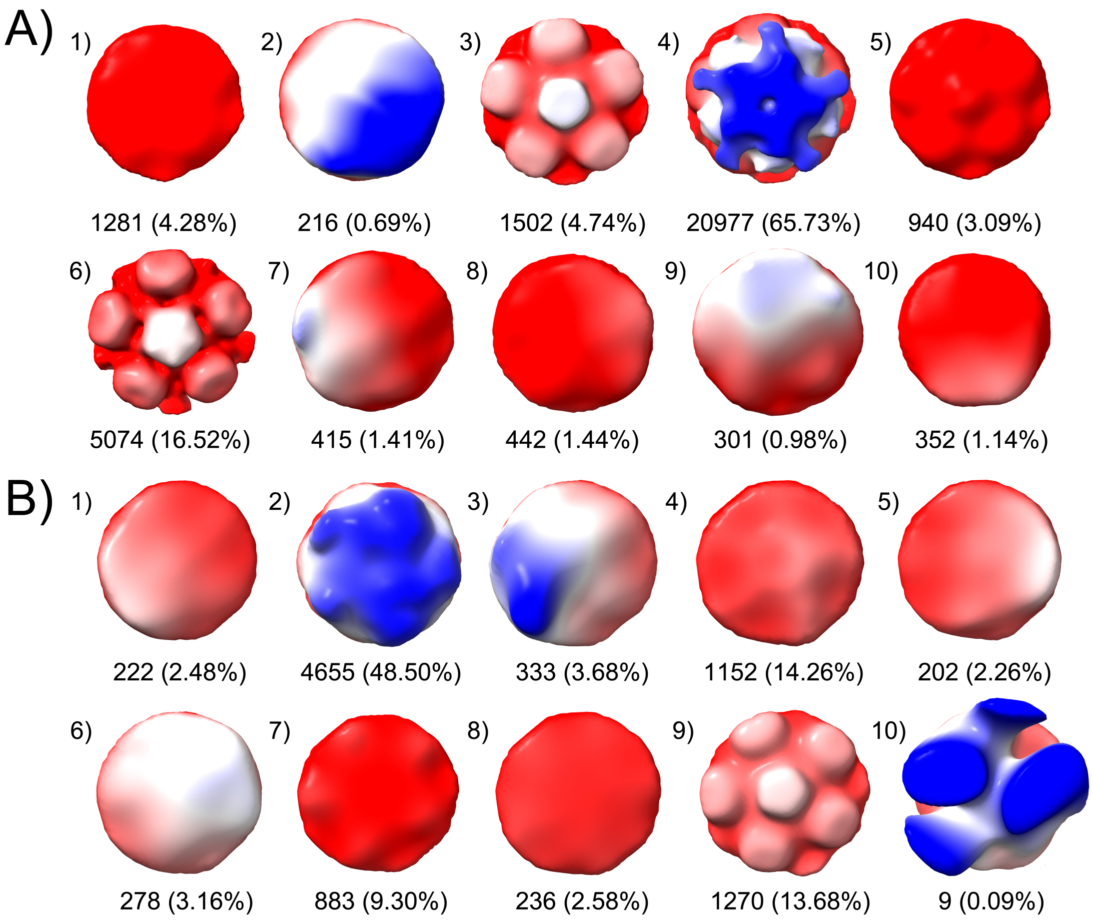


**Figure S3. Asymmetric focused classification of HSV capsids.**

Focused classification over a single 5-fold penton vertex with a T number of 5 was performed on A, B and C capsids to obtain structural information pertaining to any asymmetric features on the penton-vertices. The 10 classes calculated along with percentage of particles representing each of the classes is shown for both **(A)** B-capsids and **(B)** C-capsids. While A-capsids did not show any distinct density among any of the classes (data not shown), 1 identifiable class for both B-capsids (class 4) and C-capsids (class 2) was observed corresponding to visible pentaskelion density pertaining to CATC.

**Supplemental Video 4. Tomograms of wild type HSV capsid within the cell cytoplasm and nucleus.**

Movie showing serial sections through the z-axis of the tomogram of HSV capsids located via correlative imaging within the cytoplasm and the nucleus. Movie created using IMOD and Quicktime.

**Supplemental Video 5. Tomograms of UL37-null mutant HSV capsids within the nucleus.**

Movies denoting serial sections through tomograms of the UL37 mutant HSV capsids within the nucleus of a cell. Capsids are well dispersed throughout the nucleus and the three different capsids namely, A-, B- and C-type can be seen clearly. Movies were created using IMOD and Quicktime.

**Supplemental Video 6. *In situ* 3D reconstructions of intranuclear HSV capsids - A, B and C.**

Movies depicting the 3D reconstructions of the subclasses of capsids (A, B and C) within the nucleus obtained by subtomogram averaging. Variation in the additional density over the penton corresponding to the CATC can be clearly seen, with the C-capsids exhibiting the most apparently visible star-shaped density. Movies were created using ChimeraX.
